# NMD-mediated control of Tor influences adaptation to nutrient and temperature conditions in *Cryptococcus neoformans*

**DOI:** 10.64898/2026.02.19.706931

**Authors:** Sean R. Duffy, John C. Panepinto

## Abstract

The yeast *Cryptococcus neoformans* is an opportunistic human pathogen capable of surviving within various environmental conditions. The repertoire of antifungal agents effective in treating cryptococcal infection is limited, necessitating the identification of alternative treatment strategies. Nonsense-mediated decay (NMD) is an RNA decay mechanism that serves as a post-transcriptional regulator of gene expression. While the absence of NMD in *C. neoformans* sensitizes cells to the antifungal fluconazole, the mechanism underlying this sensitivity and role of NMD in *C. neoformans* biology remained unexplored. Using phenotypic analysis and RNA-sequencing analysis, we identify basal dysregulation of thermal- and nutrient-adaptive genes and demonstrate temperature- and/or nutrient-dependent phenotypic suppression of *upf1*Δ phenotypes, including fluconazole sensitivity and resistance to rapamycin. We determine rapamycin co-treatment also suppresses the *upf1*Δ fluconazole sensitivity, implicating dysregulation of Tor signaling in phenotypic outcomes when NMD is absent. We then investigate Tor-sensitive signaling in the *upf1*Δ mutant, finding inhibition of cell wall integrity (CWI) signaling and hyperactivation of the kinase Gcn2, both of which returned to wildtype-like levels by either rapamycin treatment, nutrient limitation, or constitutive thermal stress. These results indicated NMD is required for appropriate regulation of Tor signaling in unstressed conditions and suggested *upf1*Δ phenotypes are driven in part by Tor hyperactivation. A phenotypic screen of mutants lacking Tor regulators revealed that deletion of the Tor-suppressing *IML1* gene recapitulates *upf1*Δ phenotypes and signaling defects, consistent with Tor hyperactivation. Taken together, our results suggest NMD participates in the regulation of Tor signaling in *C. neoformans*. Future work will investigate how specific targets of NMD impact Tor signaling and promote fluconazole sensitivity in *C. neoformans*.

**Importance:** Pulmonary and central nervous system infections cause by *Cryptococcus neoformans* are responsible for about 112,000 deaths annually. Ten-week mortality remains high at 25% with use of frontline antifungals which imposes major health risks due to inherent toxicity. Thus, a need arises to identify novel avenues of treatment, including ways of boosting the efficacy of widely available antifungals such as fluconazole against C. neoformans. The design of NMD inhibitors is an active pharmaceutical pipeline for use in treating human genetic diseases. Even though NMD is conserved across eukaryotes, underlying components and regulatory roles of NMD differ between humans and fungi. Therefore, understanding NMD within *C. neoformans* will inform the design and repurposing of NMD inhibitors to enhance the antifungal activity of fluconazole as a treatment for Cryptococcosis.

## Introduction

*Cryptococcus neoformans* is a basidiomycete yeast and opportunistic human pathogen with ubiquitous global distribution (1). Disease caused by *C. neoformans* is responsible for around 112,000 deaths annually, primarily within regions with limited access to frontline antifungals such as eastern and southern Africa (2). Cryptococcal infections, typically in individuals who are immunocompromised, can cause severe pneumonia and, following dissemination to the brain, life-threatening meningitis if left untreated (3). Even with antifungal intervention 10-week mortality from cryptococcal meningitis remains relatively high at 25-30% (4, 5), along with potential liver and kidney damage from antifungal toxicity (6). Thus, a need arises for the identification of novel cryptococcal treatment strategies that are both safe and efficacious.

Treatment with the low toxicity antifungal fluconazole is commonly used in place of or alongside frontline treatment of *C. neoformans* (4, 7). Like other triazole-class antifungals, fluconazole blocks the synthesis of the fungal sterol ergosterol by inhibiting Erg11, a lanosterol 14-alpha demethylase, leading to compromised cell integrity and toxic sterol-intermediate accumulation (8). However, fluconazole only has a fungistatic effect against *C. neoformans* and the development of heteroresistance is common in response to fluconazole monotherapy (9, 10). Investigations have led to the discovery of synthetic drugs that can potentiate fluconazole against fungal pathogens (11), including the antidepressant sertraline that increases the efficacy of fluconazole against *C. neoformans* both *in vitro* and in murine infection models (12). Despite this, sertraline was found to be ineffective in improving treatment outcomes in patients with cryptococcal meningitis (13). Thus, a need arises to further elucidate the mechanisms governing fluconazole tolerance in *C. neoformans* within the context of the human host.

*C. neoformans*’ survival within the host requires adaptation to the stressors therein. This includes oxidative stress produced by host macrophages and thermal stress from mammalian endothermy. Recent work has also implicated nutrient availability within the host in pathogenicity and antifungal susceptibility of *C. neoformans* (14). For example, glucose present in the cerebrospinal fluid drives tolerance to amphotericin B in murine models of infection (15). Additionally, caloric restriction and nitrogen source availability increase *C. neoformans*’ tolerance to fluconazole and flucytosine, respectively (16, 17). Finally, alterations in nutrient signaling pathways can alter signaling through major stress response pathways leading to changes in temperature and antifungal sensitivities as well as pathogenic capacity (18–20). Despite these observations, understanding of the mechanisms involved in coordinating stress and nutrient signaling responses is limited.

Nonsense mediated decay (NMD) is a conserved eukaryotic RNA decay pathway involved in stress adaptation in both humans and fungi (21–25). NMD acts by targeting mRNA transcripts with premature termination codons (PTCs) for degradation, canonically serving as a quality control mechanism that degrades mRNAs bearing nonsense mutations in coding sequences introduced by mutations and transcriptional errors (26). In addition, NMD can control the expression of mRNAs with unique regulatory regions such as upstream open reading frames (uORFs), alternative splice sites, ribosomal frame shifts, or alternative transcriptional start sites (LUTIs) (27).

Changes in splicing, transcription start sites, and uORF utilization occur in *C. neoformans* under various environmental conditions with the affected transcripts being subject to regulation by NMD (28–30). The function of NMD is carried out by a conserved set of upstream frameshift (Upf) proteins, with the ATP-dependent RNA helicase Upf1 being necessary for NMD in humans, fungi, and plants (31–33). In *C. neoformans*, deletion of Upf1 increases sensitivity to fluconazole and to a minor extent growth 37°C (28). This suggests that targeted inhibition of NMD in *C. neoformans* may be a viable strategy for improving the efficacy of fluconazole. The development of NMD inhibitors is an active area of pharmaceutical research for use in treating rare genetic disorders originating from nonsense mutation (34). However, the mechanism by which loss of NMD sensitizes *C. neoformans* to fluconazole has not been determined.

In this work we investigated the physiological role of NMD in *C. neoformans*. Using a *upf1*Δ mutant, we identify several novel phenotypes in the absence of NMD, including increased sensitivity to reactive oxygen species (ROS), temperature-dependent resistance to ER and cell wall stressors, and alterations in virulence factor elaboration. We also determined that NMD in *C. neoformans* participates in the regulation of genes involved in the response to thermal stress and the utilization of alternative carbon and nitrogen sources. Dysregulation of these nutrient-utilizing genes is linked to the fluconazole phenotype of the *upf1*Δ mutant, through the hyperactivation of the target of rapamycin (Tor) pathway. We determine that hyperactivation of Tor resulting from the absence of NMD is nutrient and temperature conditions, as substitution of glucose as a carbon source or constitutive growth 37°C impacted stress sensitivities and restored signaling defects in the *upf1*Δ mutant. Together, our data demonstrates the importance of NMD in *C. neoformans* as regulator of stress adaptation through modulation of Tor signaling in response to environmental nutrient and temperature conditions.

## Results

### Loss of Upf1 in *C. neoformans* increases ROS sensitivity and causes temperature-dependent resistance to tunicamycin and caspofungin

To begin investigation of the physiological role of NMD in *C. neoformans*, we first began by performing a comprehensive phenotypic analysis. WT, *upf1*Δ mutant, and *upf1*Δ::*MYC-UPF1* complement strains were subjected to serial dilution spot plate assays in the presence of various compounds and stressors. We confirmed that our *upf1*Δ mutant was sensitive to fluconazole and the insensitive to growth at 37°C as previously reported (28). We did, however, observe a slight growth defect in the *upf1*Δ mutant at 38°C (Figure 1A). Novel phenotypes we identified in the *upf1*Δ mutant includes sensitivity to exogenous reactive oxygen species (ROS) in the form of hydrogen peroxide and antimycin A, a generator of endogenous ROS (35). Altered tolerance to ROS in the absence of Upf1 is known to occur in other fungal species (23, 24). The most striking phenotype observed was that the *upf1*Δ mutant was sensitive to the ER stress inducer tunicamycin and cell wall-targeting antifungal drug caspofungin at 30°C but was resistant to the same concentration of both these drugs at 37°C (Figure 1B). This temperature-dependent phenotypic reversal was not observed with fluconazole, hydrogen peroxide, or antimycin A, which appeared more penetrant at 37°C. These results showed that, as in other fungi, NMD in *C. neoformans* is involved in mediating tolerance to oxidative stress. Additionally, the temperature-dependent nature of the *upf1*Δ mutant’s tunicamycin and caspofungin phenotypes and sensitivity to 38°C indicated a connection between NMD and thermal stress in *C. neoformans*.

**Figure 1.**
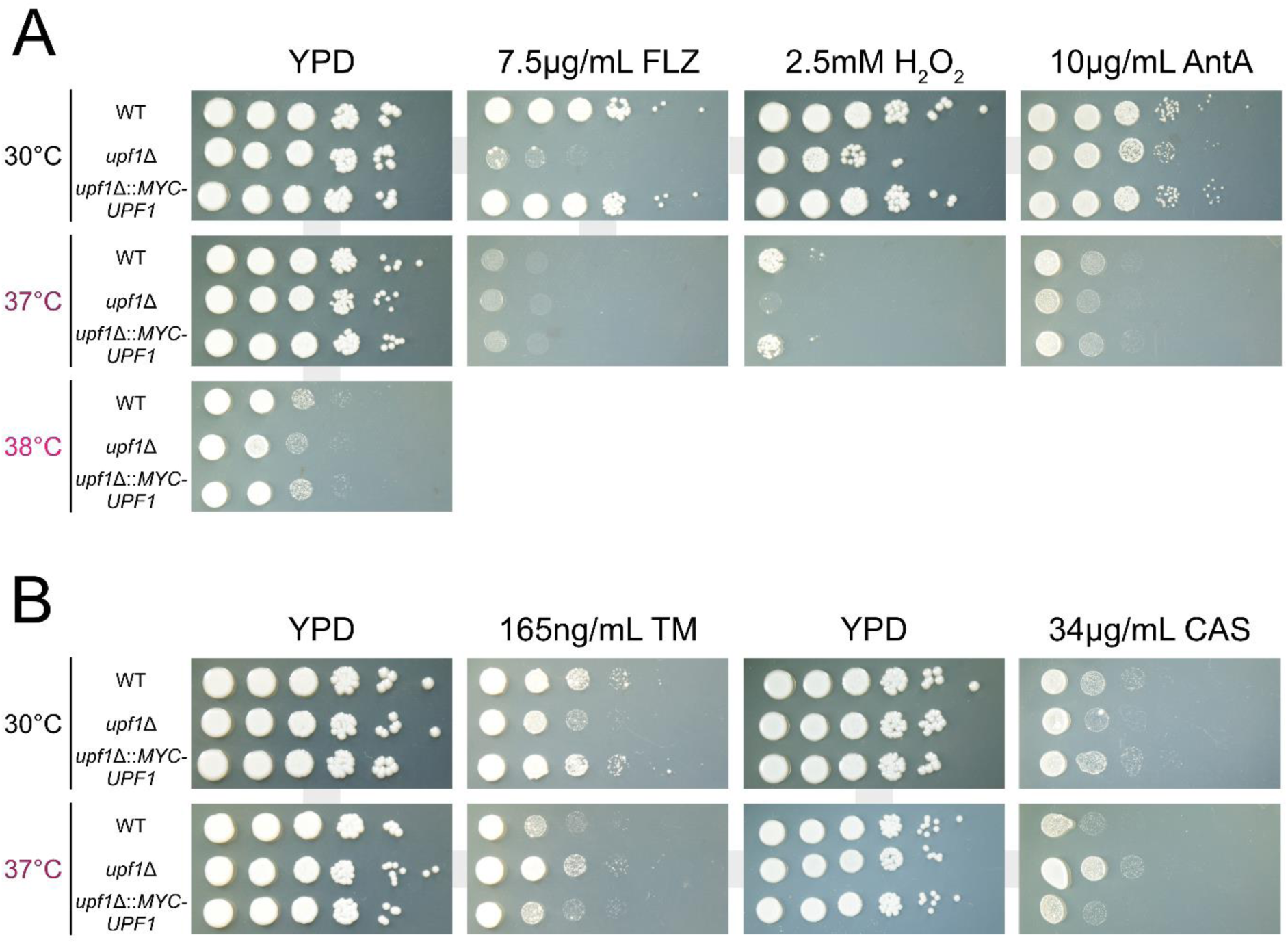
NMD is required for tolerance to reactive oxygen species and mediates a temperature-dependent response to tunicamycin and caspofungin. Serial-dilution spot plates of WT, *upf1*Δ, and *upf1*Δ::*MYC-UPF1* strains grown on yeast extract-peptone agar with 2% glucose (YPD). Drugs and compounds were added to liquid agar prior to its solidification at the indicated concentrations, including (A) fluconazole (FLZ), hydrogen peroxide (H_2_O_2_), antimycin A (AntA), (B) tunicamycin (TM), and capsofungin (CAS). Plates labelled ‘YPD’ indicate assay-specific controls where no compounds were added. Once spotted plates were incubated at the indicated temperatures for three days. Photographs in panels A and B represent separate assays performed in triplicate (n=3).

### Genes involved in nutrient utilization and thermal stress are dysregulated in the *upf1*Δ mutant

To identify the impact of NMD on transcript abundance basally and during thermal stress, RNA-seq analysis was performed on RNA samples isolated from WT and *upf1*Δ strains following growth to mid-log phase at 30°C and 1 hour after a shift to 37°C. We identified 687 significantly upregulated and 189 significantly downregulated protein-coding transcripts in the *upf1*Δ mutant compared to WT at 30°C (Data S1). We focused our analysis on genes significantly upregulated in the *upf1*Δ mutant compared to WT under both conditions as NMD-regulated transcripts are typically de-repressed in the absence of NMD. Gene ontology (GO) and pathway (KEGG) analysis of upregulated transcripts in the *upf1*Δ mutant at 30°C identified only sparse enrichment of related genes without particular significance besides transport (Figure 2A). However, thorough curation of this gene list revealed the presence of several genes associated with the transport and catabolism of various alternative carbon and nitrogen sources (Table S2). Interestingly, in this set of upregulated genes were those under forms of catabolite-mediated control in both *C. neoformans* and other fungi, including the galactose (*GAL1*, *GAL7*, and *UGE2*) and inositol (*INO1*, *MIO1*, *MIO2,* and *MIO3*) metabolic regulons (36–39). Upregulation of these genes in the *upf1*Δ mutant despite growth in nutrient-rich media (YPD), including the preferred carbon source glucose, indicated that NMD may regulate nutrient utilization from a post-transcriptional level.

**Figure 2.**
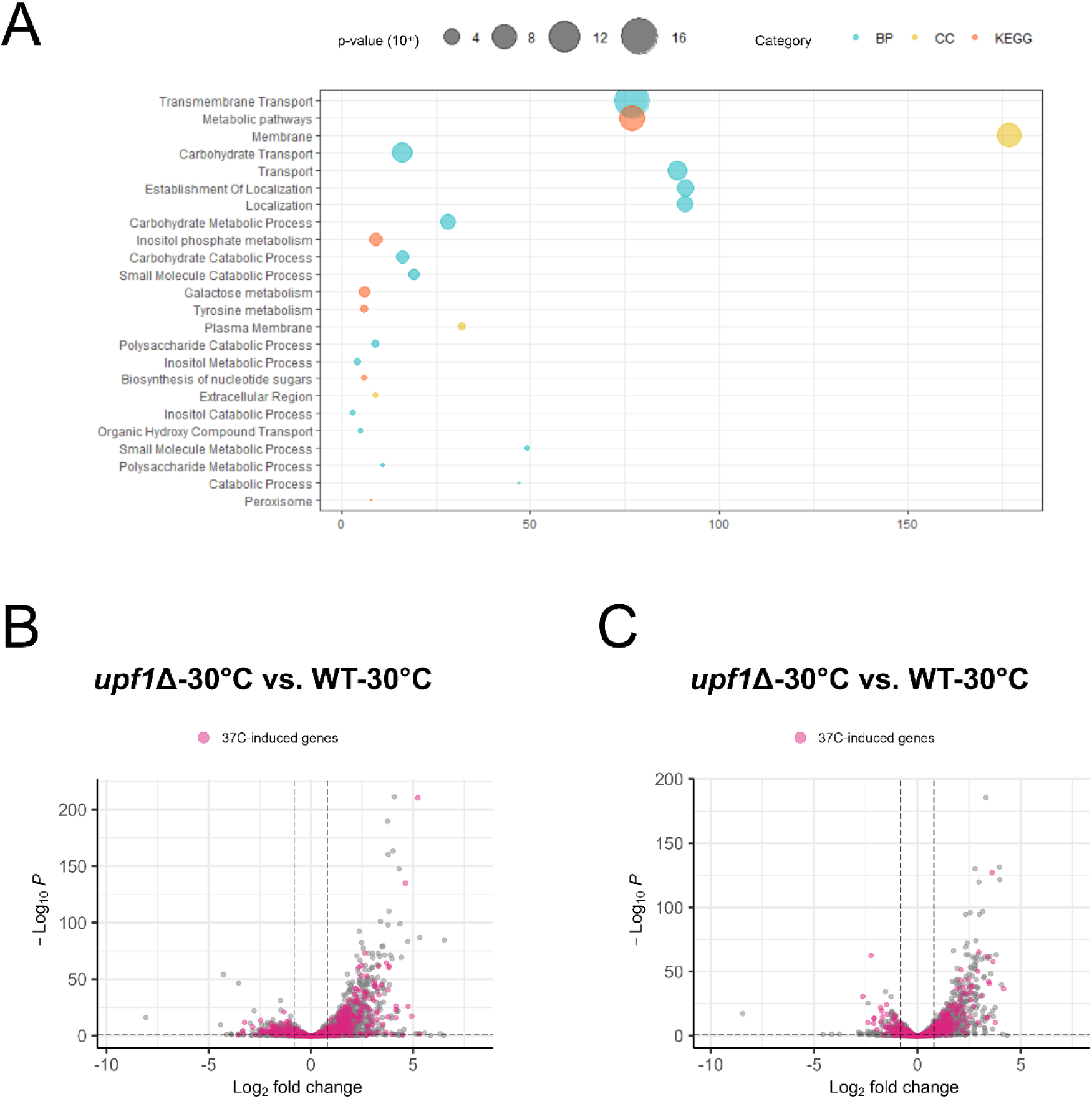
Expression of nutrient utilization and temperature-responsive genes is increased in the absence of NMD. A) Bubble plot representing results from the DAVID webserver functional annotation analysis of upregulated genes in the *upf1*Δ mutant at 30°C. The x-axis indicates the number of genes found associated with their given annotation on the y-axis. The size of each point represents the calculated p-value (x=10^-x^) for each annotation while color indicates terms for which annotation represent (blue = biological process (BP), yellow = cellular compartment (CC), orange = KEGG pathway (KEGG)). Volcano plots depict the directionality of 37°C-induced genes in the compared expression profiles of *upf1*Δ vs. WT at either (B) 30°C mid-log or (C) 37°C for 1 hour. 37°C-induced genes are defined as having log_2_(fold-change) ≥ 0.8 and adjusted p-value ≤ 0.05 in the WT, 37°C vs. WT, 30°C dataset.

**Figure 3.**
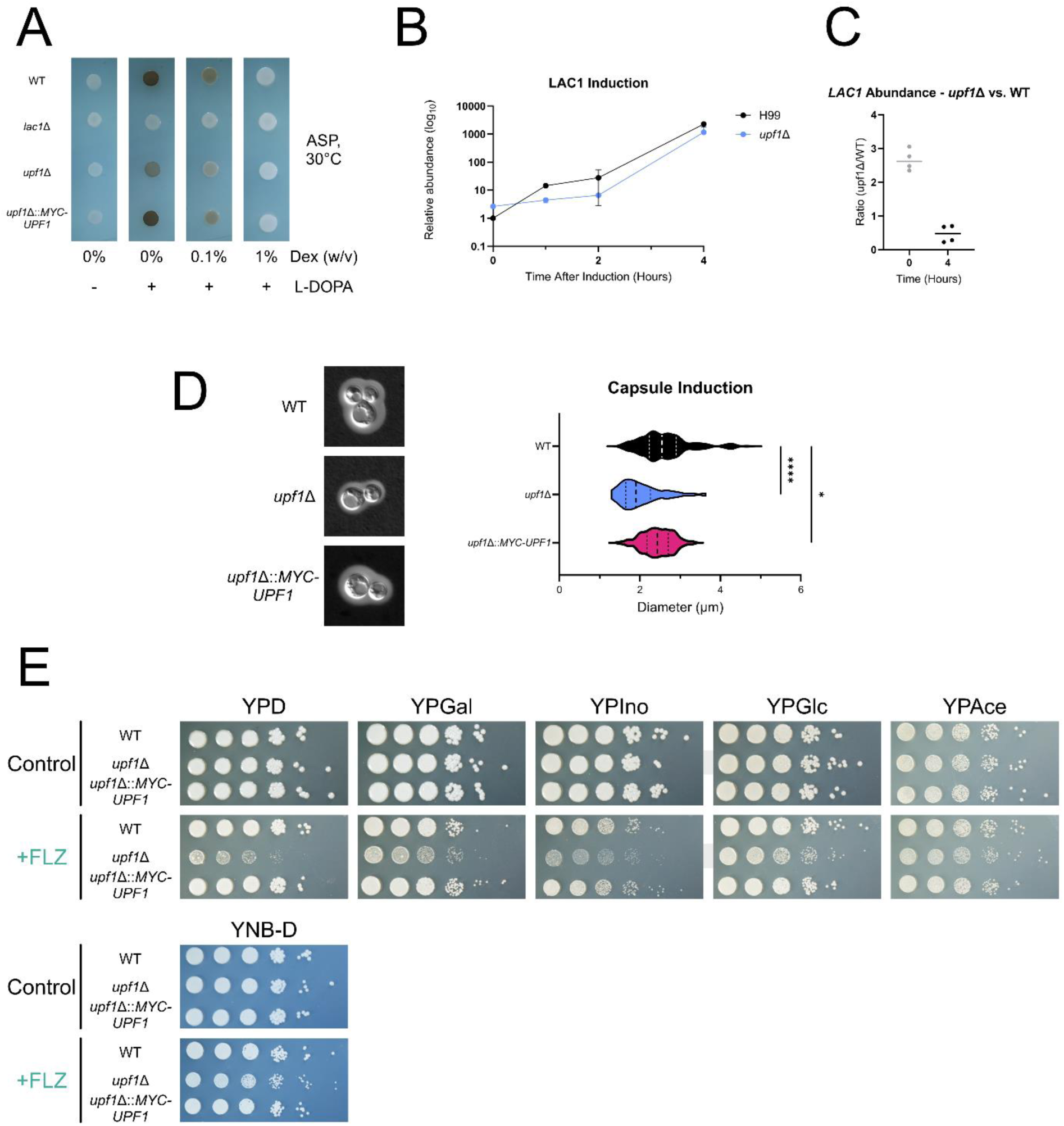
The absence of NMD attenuates virulence factor elaboration and causes nutrient source-dependent fluconazole sensitivity. A) Spot plates of WT, *lac1*Δ, *upf1*Δ, and *upf1*Δ::*MYC-UPF1* strains on ASP agar supplemented with (+) or without (-) L-dopamine (L-DOPA, 1mM) and glucose (Dex, % w/v). The *lac1*Δ mutant served as our a-melanistic control in each plate. Spotted plates were incubated at 30°C for 24 hours before being photographed. Images represent photos from same biological replicate (n=3). B-C) Relative abundance of *LAC1* mRNA measured by RT-qPCR analysis in WT and *upf1*Δ strains following a shift from YPD to ASP media. Samples were harvested from initial mid-log phase cultures (hour-0) as well as 1,2, and 4 hours post-shift to ASP. Relative abundance of *LAC1* was determined using the relative standard curve method with *GPD1* serving as the control. Graphs depict values from two technical replicates across 2 biological replicates (t = 2, n=2). D) Representative images and measurements of capsule thickness (micrometer, µm) from WT, *upf1*Δ, and *upf1*Δ::*MYC-UPF1* strains grown in capsule inducing conditions (0.1xSD media, 37°C). Graph represents the measurements of 50 cells across two biological replicates (n=2). E) Serial-dilution spot plates of WT, *upf1*Δ, and *upf1*Δ::*MYC-UPF1* strains grown in the presence or absence of fluconazole (FLZ; 7.5µg/mL) on yeast-peptone (YP) agar plates supplemented with 2% (w/v) of either glucose (YPD), galactose (YPGal), inositol (YPIno), glycerol (YPGlc), or sodium acetate (YPAce). Strains were also grown on plates containing yeast nitrogen base agar with 2% (w/v) glucose in the presence and absence of fluconazole. Photos represent plates from the same biological replicate (n=3).

**Figure 4.**
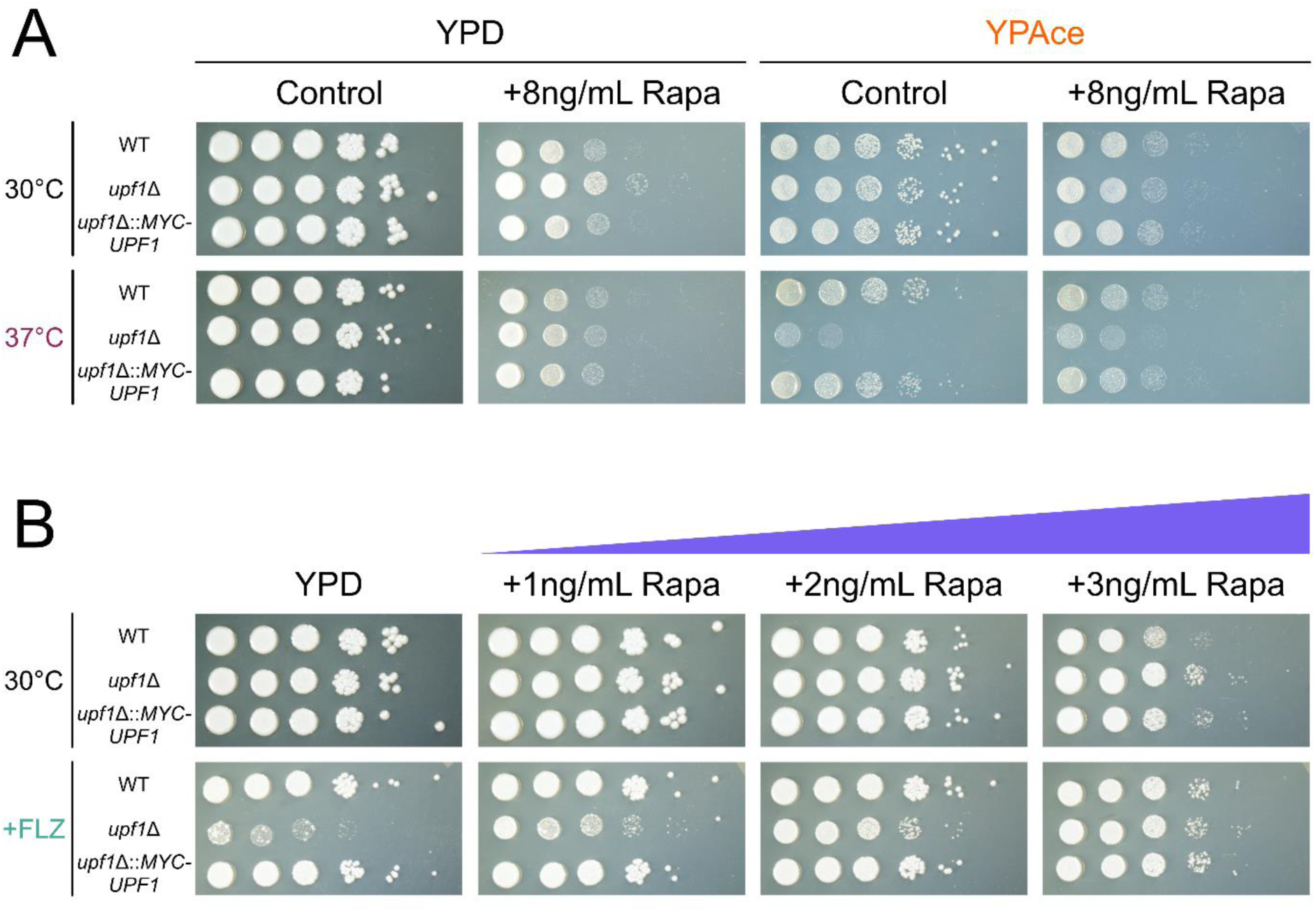
Nutrient- and temperature-sensitive changes in Tor underly rapamycin resistance and fluconazole sensitivity of the *upf1*Δ mutant. A) Serial-dilution spot plates of WT, *upf1*Δ, and *upf1*Δ::*MYC-UPF1* strains grown on YPD or YPAce agar supplemented with or without rapamycin (Rapa; 8ng/mL). Duplicate spot plates for each condition were made and split between incubation at 30°C or 37°C. B) Serial-dilution spot plates of WT, *upf1*Δ, and *upf1*Δ::*MYC-UPF1* strains where YPD agar with or without the addition of fluconazole (FLZ; 7.5µg/mL) were supplemented with increasing dosages of rapamycin (Rapa; 1,2 or 3ng/mL). Photos represent plates from the same biological replicate (n=3) after three days of incubation.

Comparison of *upf1*Δ mutant and WT 37°C-Shift conditions revealed 624 and 184 significantly upregulated and downregulated transcripts, respectively. To investigate possible connections between NMD and thermal stress, we compared the expression of genes in the *upf1*Δ mutant to the set of gene differentially expressed in WT in response to thermal stress. Transcripts differentially expressed by WT in response to 37°C were highlighted within volcano plots of *upf1*Δ and WT at 30°C and 37°C. We found that transcripts upregulated in the *upf1*Δ mutant at 30°C were enriched in genes upregulated in response to 37°C in the wildtype (Figure 2B). Additionally, these 37°C-induced transcripts were also enriched in the comparison of the *upf1*Δ mutant and WT at 37°C (Figure 2C). Together, these data indicate that NMD in *C. neoformans* represses a subset of temperature-responsive transcripts whose increased expression during thermal stress partially relies on escape from or changes in NMD efficacy.

### Contradictory responses to environmental conditions drive virulence factor attenuation and nutrient-dependent fluconazole sensitivity in the *upf1*Δ mutant

The transcriptional profile of the *upf1*Δ mutant led us to hypothesize that NMD may regulate other processes influenced by temperature and nutrients in *C. neoformans*, such as virulence factor elaboration. The induction of the virulence factor melanin in *C. neoformans* is known to require starvation-like conditions (40). In our RNA-seq, the transcripts encoding the melanin-synthesizing laccases (*LAC1* and *LAC2*) was significantly upregulated in the *upf1*Δ mutant (Data S1). Therefore, we performed spot plate assays on a melanin-inducing medium with L-DOPA to determine relative levels of pigmentation between WT, *upf1*Δ, *upf1*Δ::*MYC-UPF1*, and *lac1*Δ strains.

Surprisingly, the *upf1*Δ mutant showed reduced pigmentation compared to either the WT or complement strains, despite the apparent increase in laccase mRNA expression (Figure 2A). Using RT-qPCR analysis, we found that the *upf1*Δ mutant had reduced *LAC1* mRNA levels compared to WT following a shift melanin-inducing medium, despite having a higher initial abundance (Figure 2B-C). Induction of the antiphagocytic capsule in *C. neoformans* is complex, including inputs from nutrient limitation and temperature (41). The myo-inositol oxygenase proteins Mio1-3, whose transcripts we found upregulated in the *upf1*Δ mutant, promote the formation of capsule (39). Thus, capsule thicknesses were measured for WT, *upf1*Δ, and *upf1*Δ::*MYC-UPF1* strains grown in a capsule-inducing conditions (10% Sabouraud-dextrose media, 37°C) following India ink staining. We were again surprised to see that the *upf1*Δ mutant had a reduced capsule size than WT and complement strains (Figure 2D). These results suggested a seemingly contradictory relationship between the transcriptional profile and phenotypic outcomes of the *upf1*Δ mutant. We wonder how other phenotypes may result arise from opposing directionality of gene regulation in the *upf1*Δ mutant, particularly the *upf1*Δ mutant’s sensitivity to the antifungal fluconazole. Growth in calorically restrictive low-glucose conditions has been shown to increase fluconazole tolerance in *C. neoformans* (16). We hypothesized that the improper expression of nutrient utilization genes in the *upf1*Δ mutant under nutrient-rich conditions may have the opposite effect leading to a “caloric surplus” that decreases fluconazole tolerance. To test this hypothesis, we subjected WT, *upf1*Δ, and *upf1*Δ::*MYC-UPF1* strains to serial-dilution spot plate assays on yeast extract-peptone (YP) agar plates with-or-without fluconazole supplemented with carbon sources that require alternative routes for use in central carbon metabolism. Additionally, we included yeast nitrogen base agar (YNB-D) plates where ammonium sulfate served as the limiting source for nitrogen metabolism. We observed that, apart from inositol, growth on alternative carbon sources suppressed the fluconazole sensitivity of the *upf1*Δ mutant to varying degrees compared to glucose (Figure 2E). Suppression of the *upf1*Δ mutant’s fluconazole sensitivity was also observed when grown under conditions of nitrogen source limitation. The most pronounced restoration was observed from growth on acetate, a nonfermentive carbon source that is directly converted into acetyl-CoA for use by cellular metabolism (42). These results indicated that the fluconazole phenotype of the *upf1*Δ mutant is linked to an altered response to nutrient availability.

### Altered signaling through the target of rapamycin (Tor) pathway underlies the fluconazole phenotype of the *upf1*Δ mutant

If the upregulated transcripts in *upf1*Δ mutant are resulting in a nutrient surplus, pathways involved in sensing cellular nutrient conditions may also be dysregulated. One pathway implicated in the response to calorically restricted conditions in *C. neoformans* (43) and intersects with NMD in other model organisms is the target of rapamycin (Tor) pathway. Tor is highly conserved eukaryotic signaling network that controls growth, catabolism, and cellular dynamics with respect to nutrient availability (44). Regulatory links between Tor and NMD have also been identified in humans and fungi (45–48). Tor in *C. neoformans* regulates growth in response to temperatures stress and the presence of fluconazole (49, 50). Thus, to investigate whether NMD and Tor are connected in *C. neoformans*, we first compared the sensitivity of WT, *upf1*Δ, and *upf1*Δ::*MYC-UPF1* strains to the Tor kinase inhibitor rapamycin (51). Using serial dilution spot plate assays, we saw that the *upf1*Δ mutant was resistant to rapamycin on YPD media at 30°C. Interestingly, resistance to rapamycin was not seen in the *upf1*Δ mutant when grown at 37°C or on YPAce (Figure 5A), suggesting that the underlying mechanism of rapamycin resistance in the *upf1*Δ mutant is suppressed or relieved under these conditions. Since growth on YPAce had a similar yet inverse repressive effect on fluconazole sensitivity of the *upf1*Δ mutant, and the connection between Tor and fluconazole sensitivity in *C. neoformans* (50), we investigated if co-treatment with rapamycin would influence the fluconazole phenotype of the *upf1*Δ mutant. Our spot plate analysis showed that rapamycin co-treatment suppressed the fluconazole sensitivity of the *upf1*Δ mutant in a dose-dependent manner (Figure 5B). Taken together, these data suggest that Tor signaling is elevated in the absence of NMD in *C. neoformans*, driving rapamycin resistance and fluconazole sensitivity in the *upf1*Δ mutant. Suppression of this rapamycin resistance by YPAce or 37°C growth, as well as the suppression of the fluconazole sensitivity by rapamycin or YPAce, also suggested that the changes in Tor signaling of the *upf1*Δ mutant are still responsive to environmental conditions known to suppress Tor signaling. Indeed, changes in the expression of Tor signaling genes is observed in *C. neoformans* under low-glucose conditions (43). Additionally, overexpression of the Tor1 kinase in *C. neoformans* leads to thermal stress sensitivity, which can be reversed by rapamycin (49).

**Figure 5.**
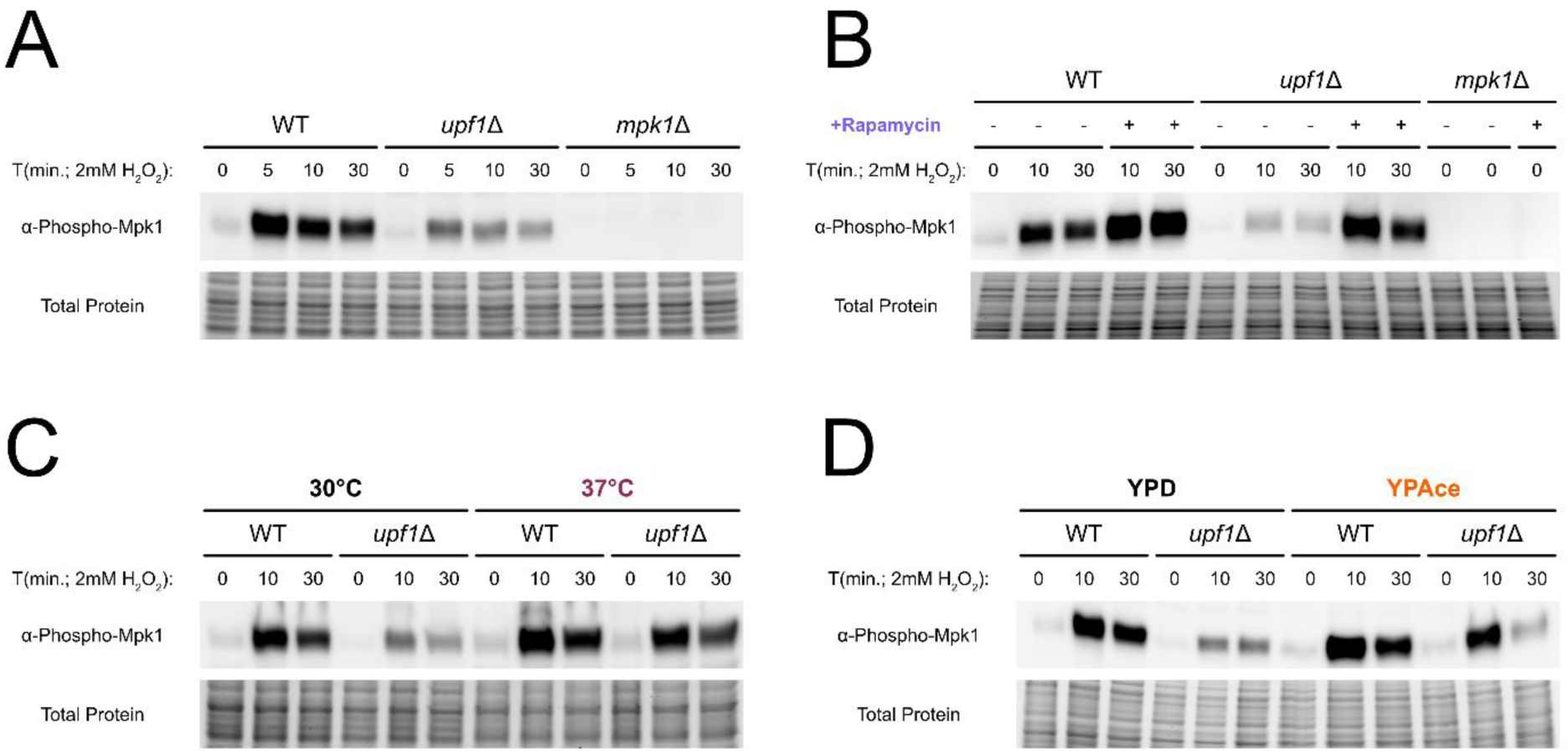
Hyperactive Tor represses CWI signaling in the *upf1*Δ mutant and is de-repressed by rapamycin, temperature and nutrients. A) Western blots for phospho-Mpk1 in WT, *upf1*Δ, and *mpk1*Δ strains in response to 2mM hydrogen peroxide (H_2_O_2_). B) For experiments assessing phospho-Mpk1 during rapamycin co-treatment, cultures were split following harvest of initial time point samples (T = 0 minutes). Then, cultures were treated with hydrogen peroxide alone (-) or with hydrogen peroxide after the addition of rapamycin (+; 1µg/mL). C-D) For experiments assessing phospho-Mpk1 under thermal and nutrient stress, control and experimental cultures for both WT and *upf1*Δ mutant were separately seeded and grown to mid-log phase prior to treatment with hydrogen peroxide. Control cultures for both experiments were grown in YPD at 30°C. For experimental cultures, only one variable per experiment was altered compared to the control culture, being either the temperature of constitutive growth (C; 30°C vs. 37°C) or the media cultures were seeded in (D; YPD vs. YPAce). Blots are representative of individual biological replicates (n=3).

### Suppression of Mpk1 activation by hyperactive Tor in the *upf1*Δ mutant can be rescued by rapamycin treatment, nutrient limitation, or thermal stress

To gain clarity on the relationship between NMD of Tor signaling in *C. neoformans*, we set out to compare flux through Tor-regulated signaling pathways in WT and the *upf1*Δ mutant. To this end, we chose to measure signaling through the Mpk1 MAP kinase, a component of the cell wall integrity (CWI) pathway in *C. neoformans* which promotes survival to thermal, cell integrity, and oxidative stress (52).

Phosphorylation of Mpk1 through the CWI pathway is negatively regulated by Tor signaling in *C. neoformans* and *S. cerevisiae* (49, 53, 54). Thus, we assessed levels of phosphorylated Mpk1 (phospho-Mpk1) by western blot analysis of protein isolated from both WT, *upf1*Δ, and *mpk1*Δ strains following treatment hydrogen peroxide, a potent inducer of the CWI pathway (55). We observed reduced induction of phospho-Mpk1 in the *upf1*Δ mutant compared to WT following CWI stimulation (Figure 6A). Indeed, treatment with rapamycin immediately prior to CWI activation was able to de-repress phospho-Mpk1 induction in the *upf1*Δ mutant (Figure 6B), supporting that CWI suppression in the *upf1*Δ mutant is Tor-mediated. Since nutrient and temperature conditions suppressed rapamycin resistance in the *upf1*Δ mutant, we investigated whether these conditions also altered signaling through the CWI. We observed de-repression of phospho-Mpk1 induction in the *upf1*Δ mutant under constitutive growth at 37°C compared to constitutive growth at 30°C (Figure 6C). Growth in YPAce was also able to de-repress phospho-Mpk1 induction in the *upf1*Δ mutant compared to YPD but did not maintain de-repression through the last time point measured (Figure 6D). This data shows that the absence of NMD in *C. neoformans* leads to Tor hyperactivation which suppresses the CWI. De-repression of the CWI in the *upf1*Δ mutant can consequently occurs though either direct inhibition of the Tor kinase or in environmental conditions that suppress Tor.

**Figure 6.**
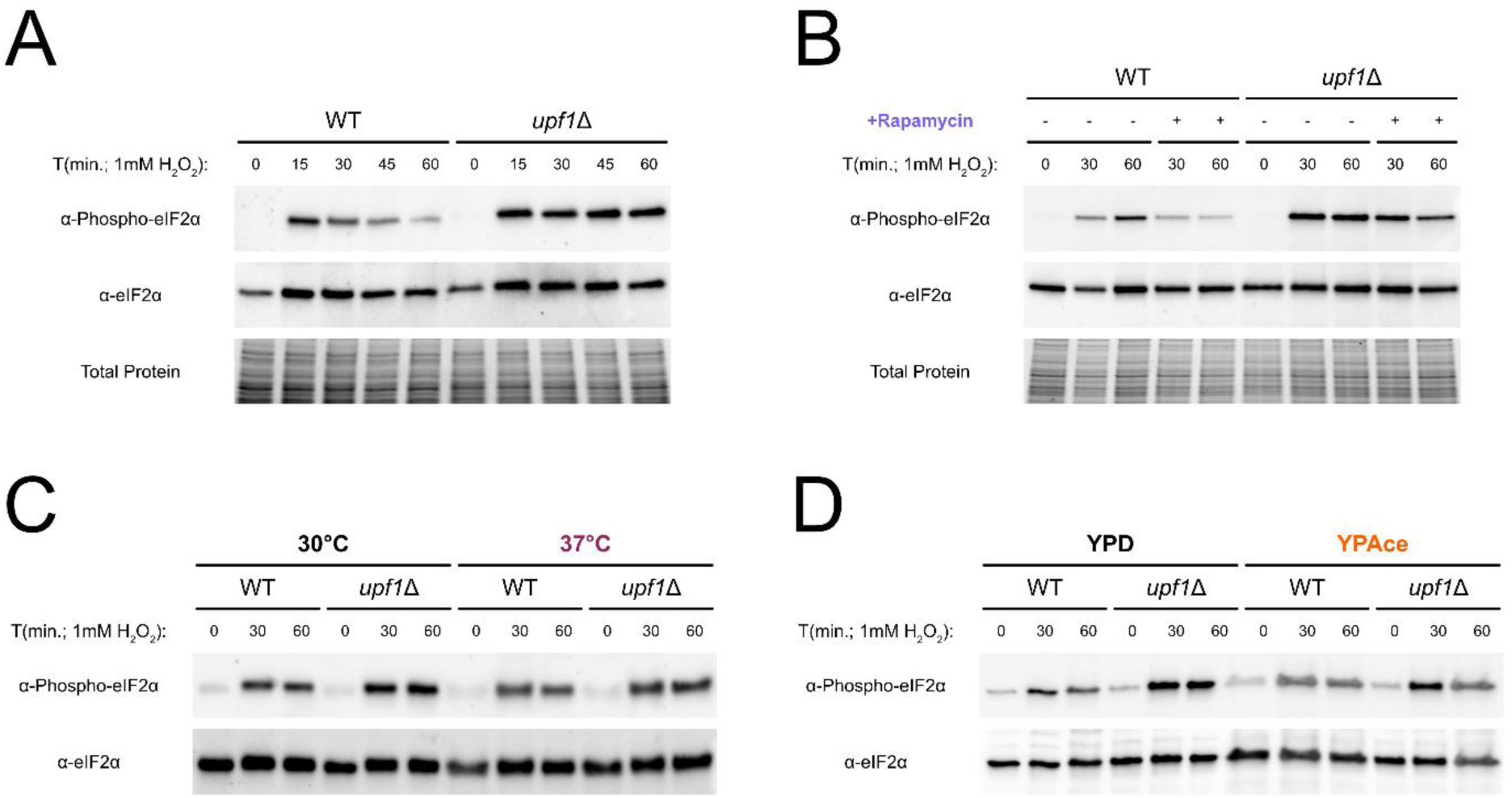
Hyperactivation of Tor elicits increased Gcn2-mediated eIF2α phosphorylation in the *upf1*Δ mutant. A-B) Western blots for phospho-eIF2α in WT and *upf1*Δ strains in response to 1mM hydrogen peroxide (H_2_O_2_). For experiments involving rapamycin co-treatment (B), cultures were split prior to the start of the time course with half the culture receiving only hydrogen peroxide treatment (-) and the other receiving hydrogen peroxide treatment immediately following the addition of rapamycin (+; 1µg/mL). C-D) For assessment of phospho-eIF2α levels between temperature and nutrient conditions, a control and experimental culture for both WT and *upf1*Δ strains were used, with control cultures being grown in YPD at 30°C. The experimental cultures differed from controls either by the constitutive temperature of growth (C; 37°C) or the medium the culture (D; YPAce). Isolated samples were then used in western blotting for total levels of eIF2α. Then, densitometry of total eIF2α signal from each lane was used to determination the relative abundance per volume of each sample (AU_eIF2α_/µL). Western blot analysis was then repeated using the calculated volumes of sample necessary for equal levels of eIF2α (1*10^6^ AU_eIF2α_/well). The resulting blots were propped for phospho-eIF2α before being stripped and probed for total eIF2α (n=3).

### Increased activation of the Gcn2 kinase in the *upf1*Δ mutant indicates the presence of conserved response to Tor hyperactivation in *C. neoformans*

With limited information surrounding downstream targets of Tor in *C. neoformans*, we explored literature of other systems to identify any conserved consequences of Tor hyperactivation. We identified that activation of the eIF2α kinase Gcn2, a conserved sensor of amino acid deprivation (AAD) in eukaryotes (56, 57), occurs during improper-or hyper-activation of Tor in humans and several fungi, where Gcn2 suppresses Tor (58–60). In fact, direct and indirect interactions between Gcn2 and Tor are important in humans and fungi for coordinating growth, translation, metabolism, autophagy, and stress responses with respect to environmental conditions (61–64). Our lab has identified the importance of Gcn2 in *C. neoformans* as a translational regulator with roles in nitrogen source utilization, oxidative stress, and thermal stress (65–67). Thus, Gcn2 activity was assessed by Western blot analysis measuring levels of phosphorylated eIF2α (phospho-eIF2α) in both WT and the *upf1*Δ strains following a mild treatment of hydrogen peroxide. We found that the *upf1*Δ mutant had persistent and increased levels of phospho-eIF2α post-treatment compared to WT (Figure 7A). To determine if this increase in eIF2α phosphorylation was Tor-dependent, we assessed whether rapamycin co-treatment, growth in YPAce, or growth at 37°C altered phospho-eIF2α induction in WT and the *upf1*Δ mutant. Interestingly, rapamycin co-treatment reduced phospho-eIF2α in both WT and the *upf1*Δ mutant compared to untreated controls (Figure 7B), suggesting that Tor modulates Gcn2 activation basally in *C. neoformans*. While phospho-eIF2α levels in the *upf1*Δ mutant were reduced at 37°C and in YPAce, total eIF2α levels was also decreased under these conditions (Figure S1A-B), possibly due to changes in translational capacity in response to temperature and nutrient conditions (68, 69). To assess relative changes in phospho-eIF2α levels at 37°C or in YPAce, we re-performed western blot analysis where total eIF2α served as the loading control instead of total protein. Using this approach, we determined that growth at 37°C and YPAce does repress relative levels of phospho-eIF2α induction in the *upf1*Δ mutant (Figure 7C-D). These results demonstrate that crosstalk between Gcn2 and Tor is conserved in *C. neoformans* and is responsible for the changes in eIF2α phosphorylation seen in the *upf1*Δ mutant.

**Figure 7.**
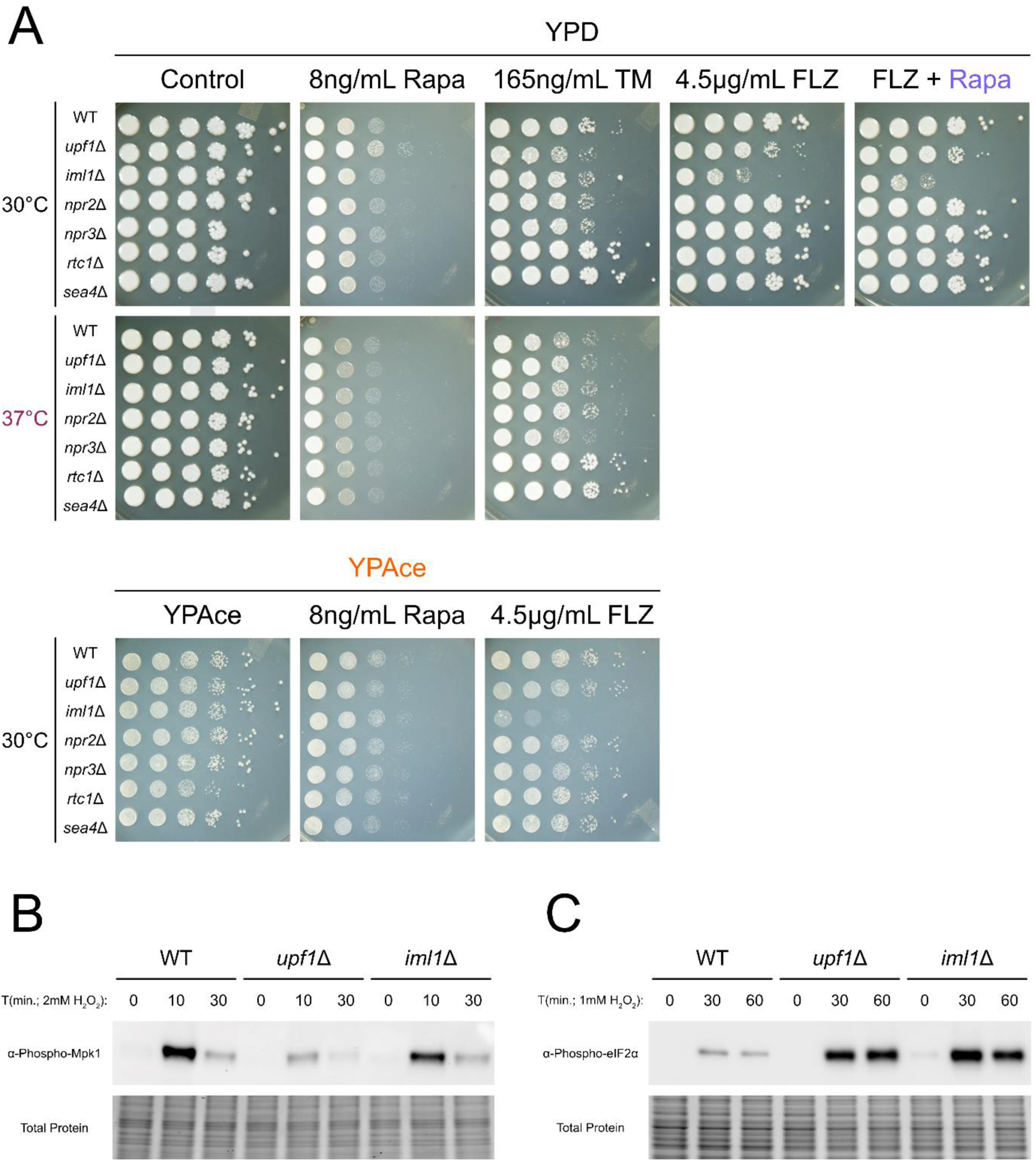
Absence of Gator1 inhibition on Tor has similar consequences on growth and signaling as the absence of NMD. A) Serial-dilution spot plates of WT, *upf1*Δ, *iml1*Δ, *npr2*Δ, *npr3*Δ, *rtc1*Δ, and *sea4*Δ KN99 strains under conditions that were identified to affect the H99 *upf1*Δ mutant. A reduced concentration of fluconazole (4.5µg/mL) was used due to inherent increased sensitivity to fluconazole of KN99 strains. This concentration was also used in the presence of rapamycin (FLZ + Rapa; Rapa = 1.5ng/mL). Western blotting for phospho-Mpk1 (B) and phsopho-eIF2α (C) in WT, *upf1*Δ, and *iml1*Δ strains following treatment with hydrogen peroxide. Blots are representative of individual biological replicates (n=3).

### Mutants lacking components of the Tor-inhibiting Gator1 complex share phenotypes and signaling defects with the *upf1*Δ mutant

Our data thus far suggested that hyperactivation of Tor in the *upf1*Δ mutant likely originates from increased sensing of nutrients. We reasoned that deletion of upstream nutrient-dependent regulators of Tor may therefore result in phenotypes like those observed in the *upf1*Δ mutant. The Gator1 and Gator2 complexes serve as major relays of intracellular nutrient conditions to Tor in both humans and fungi, where Gator1 inhibits Tor signaling under low-nutrient conditions while Gator2 inhibits Gator1 under nutrient replete conditions (70, 71). To determine the phenotypes associated with the absence of Tor regulators, we identified the homologous Gator components present in *C. neoformans* (Table S3) and then acquired deletion mutant strains that were available (Gator1: *iml1*Δ, *npr2*Δ, *npr3*Δ; Gator2: *rtc1*Δ, *sea4*Δ) along with the *upf1*Δ mutant from the Madhani deletion collection in the KN99 background of *C. neoformans* (72). Spot plates analysis revealed that although none of the Gator mutants exhibited altered rapamycin, the *iml1*Δ mutant exhibited a greater sensitivity to fluconazole than the *upf1*Δ mutant. Interestingly, the fluconazole sensitivity of the *iml1*Δ mutant was not suppressible by rapamycin co-treatment nor growth on YPAce, unlike that of the *upf1*Δ mutant (Figure 8A). All three of the Gator1 mutants had a similar tunicamycin sensitivity to the *upf1*Δ mutant at 30°C and, interestingly, the *iml1*Δ and *npr2*Δ mutants appeared to be resistant to tunicamycin at 37°C, similar to the *upf1*Δ mutant. In contrast, Gator2 mutants were tunicamycin resistant at both 30°C and 37°C. These phenotypes indicated that hyperactivation of Tor in the *upf1*Δ mutant may occur through nutrient-mediated suppression of Gator1, as the absence Iml1, the catalytic core of Gator1, results in phenotypes like the *upf1*Δ mutant. The inability of Tor-inhibiting conditions to suppress fluconazole sensitivity of the *iml1*Δ mutant may indicate that Gator1 plays a role in nutrient- and temperature-dependent Tor inhibition. To investigate whether the absence of Iml1 affects Tor-dependent signaling in *C. neoformans*, we performed western blot analysis of phospho-Mpk1 and phospho-eIF2α following activation of the CWI and Gcn2, respectively, in KN99 WT, *upf1*Δ, and *iml1*Δ strains. Phospho-Mpk1 levels were reduced in the *iml1*Δ mutant compared to WT but not to the extent seen in the *upf1*Δ mutant (Figure 8B). However, the *iml1*Δ mutant did exhibit increased and prolonged induction of phospho-eIF2α similar to the *upf1*Δ mutant (Figure 8C). Together, these data are consistent with our model that suppression of the Gator1 complex by increased nutrient signaling likely contributes to several of the growth and signaling phenotypes observed in the *upf1*Δ mutant. Further work is needed to tease out the dynamics underlying Gator control of Tor in *C. neoformans*, as the net effect of activating and inhibitory nutrient and environmental signals is complex and varies between organisms (73).

**Figure 8.**
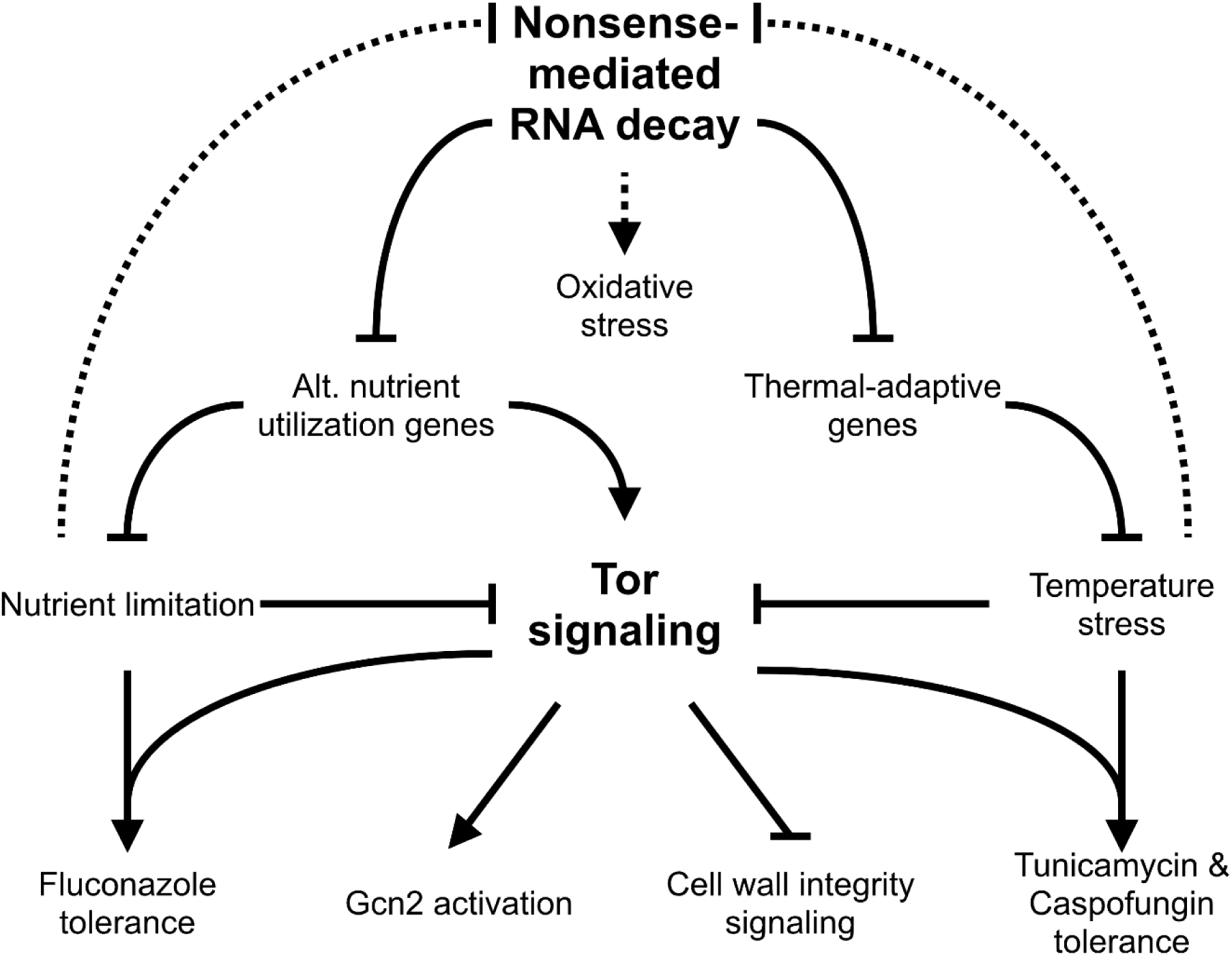
Model of NMD-mediated control of nutrient- and thermal-stress adaptation involving environmental suppression of Tor signaling. In the absence of stress NMD represses expression of alternative-nutrient utilizing and thermal adaptive genes post-transcriptionally through targeted mRNA degradation. In response to nutrient or temperature stress, signaling through Tor is suppressed, temporarily reducing growth and prompting changes in signaling and translation that promote induction of stress-adaptive regulons. Expression of NMD-targeted transcripts increases in response to stress, potentially through suppression of NMD or escape from NMD surveillance. Translation of these transcripts produces factors that combat the inhibitory nature of nutrient and thermal stress while also stimulating the re-activation of Tor. As stress is resolved, NMD re-establishes control over these stress-adaptive genes and returns their expression to a repressed state.

## Discussion

Adaptation to the stressors present within the human host microenvironment is paramount to the success of *C. neoformans* as a pathogen. Coordination between stress-responsive and environmental-sensing pathways ensures that the allocation of resources to cellular processes can be adjusted to meet the needs of the cell under a variety of environmental circumstances. Our work supports a model where NMD in *C. neoformans* regulates thermal stress adaptation and nutrient-dependent activation of Tor through post-transcriptional control of transcript abundance (Figure 9). The directionality of regulation would imply that these transcripts escape NMD regulation during thermal and nutrient stress to promote adaptation. Dysregulation of this response would explain why phenotypes of the *upf1*Δ mutant, such as the altered tolerances to fluconazole, rapamycin, tunicamycin, and capsofungin, manifest under unstressed conditions and are suppressed in the presence of nutrient and/or thermal stress. Partial de-repression of these stress-adaptive genes may also dampen signals reflective of the true levels of nutrient and thermal stress, explaining the attenuated virulence factor elaboration of the *upf1*Δ mutant.

Despite previous observations of poor overlap between NMD regulons of various fungi (24, 28, 74), our phenotypic and genetic characterization of the *upf1*Δ mutant in *C. neoformans* indicates the existence of a broader, shared physiological role of NMD amongst fungi. Specifically, NMD in fungi seems to participate in the control of metabolic flux through mechanisms tailored to the lifestyle of a given fungus. This apparent role of NMD as a regulator of metabolism may explain the high plasticity of transcripts under NMD control between fungal species, reflecting each organism’s environmental niche, growth patterns, preferred nutrients, and metabolic capacity. Early studies in *S. cerevisiae* showed the absence of Upf1 a respiratory growth defect, particularly at low temperatures (75). Later studies revealed that NMD in *S. cerevisiae* regulates the abundance of transcripts encoding proteins under catabolite-mediated regulation (*CPA1*, *OAZ1*, *DAL5*) (76–78), activate transcription of metabolic regulons (*GCN4*, *GAL4*, *ADR1*) (76, 79), and maintain homeostasis of bio-metals (*MAC1*, *FRE1*, *FRE2*) (80, 81). In the diurnal yeast *N. crassa*, the circadian rhythm is the central regulator of growth phases and metabolic flux (82). NMD regulates this circadian rhythm by targeting the mRNA of the FRQ protein, an essential component of the circadian clock mechanism (83). Additionally, the expression of cellulases in the wood-decaying fungus *Trichoderma reesei* is controlled by NMD with respect to the presence or absence of cellulose (48). Our data mirrors these findings as NMD appears to regulate unique aspects of *C. neoformans* biology including the production of capsule and melanin as well as thermotolerance. Utilization of inositol as a carbon source has been proposed as a metabolic niche of *C. neoformans*, possessing an expanded repertoire of ten inositol-transporter (*ITR*s) and three inositol-oxygenase (*MIO*s) genes compared to compared to the average set of three *ITR*s and one *MIO* in most fungi (39). Indeed, our RNA-seq showed that inositol metabolizing genes were upregulated in the *upf1*Δ mutant in *C. neoformans*, including *INO1*, all three *MIOs*, and several *ITR*s (Data S1). Interestingly, inositol was the only alternative carbon source tested not able to suppress the fluconazole sensitivity of the *upf1*Δ mutant. Further investigation into the set of genes upregulated in the *upf1*Δ mutant in *C. neoformans* and other fungal species is needed to fully understand NMD as a niche-tailored regulator of metabolism.

A shared physiological role of NMD amongst fungi appears to extend to processes peripheral to metabolism, such as oxidative stress resistance. ROS is predominantly generated as a consequence of metabolism by oxidative phosphorylation (84). We observed that the *upf1*Δ mutant in *C. neoformans* was more sensitive to both endogenous and exogenous sources of ROS. Mutants of *S. pombe* and *N. crassa* lacking Upf1 also exhibit altered ROS tolerance, but differ in their directionality (sensitivity and resistance, respectively) and underlying mechanism (destabilization transcription factor *ATF1* mRNA and overexpression of the catalase gene *cat-3*, respectively) (23, 24). We did not observe changes in changes expression of the homologs of *ATF1* nor *cat-3* in the *C. neoformans* upf*1*Δ mutant, suggesting that NMD regulates oxidative stress tolerance of *C. neoformans* by a distinct mechanism.

We also identified the presence of conserved crosstalk between Gcn2 and Tor in *C. neoformans*. This crosstalk manifests in the *upf1*Δ mutant as increased activation of Gcn2 following stress, which can be suppressed in conditions we found also suppress Tor. In *S. cerevisiae*, Gcn2 directly phosphorylates the TORC1 complex component Kog1 (62). This allows cells to resolve conflicts that arise when conditions for growth seem generally favorable but lack specific nutrients or nutrients in forms usable for cellular processes (59, 61). Reciprocally, Activation of Gcn2 by Tor, either directly in humans (58) or indirectly in *S. cerevisiae* (63), maintains active surveillance as intracellular nutrients content depletes to fuel the growth of the cell. Interestingly, hyperactivation of Gcn2 in the *upf1*Δ mutant may be an attempt to re-establish control of NMD. Hypoxia and ER stress induces phosphorylation of eIF2α in human cells that is inhibits NMD, de-repressing NMD-targeted transcripts that encode proteins that alleviate hypoxic- and ER-stress (21, 85). Additionally, de-repression of *cat-3* mRNA by NMD in *N. crassa* during oxidative stress requires degradation of Upf1 by an eIF2α phosphorylation-dependent mechanism (24). As the sole eIF2α kinase in *C. neoformans* (66), future investigations will determine whether Gcn2 mediates similar forms of regulation on cryptococcal NMD.

The *upf1*Δ mutant’s sensitivity to fluconazole indicates a potential route of boosting the efficacy of fluconazole in treating cryptococcal infection through the repurposing NMD inhibitors designed to treat genetic diseases in humans (34). However, the connection between NMD and fluconazole sensitivity in *C. neoformans* was unclear. Our work contextualizes the fluconazole phenotype of the *C. neoformans upf1*Δ mutant, originating nutrient-dependent hyperactivation of Tor signaling. Our analysis of mutants lacking components of Gator complexes indicates that hyperactivation of Tor may also drive the temperature-dependent tunicamycin phenotype of the *upf1*Δ mutant. Sensitivity to both tunicamycin and caspofungin have been reported in an *S. cerevisiae* strain with an allele that hyperactivates the Tor kinase (54). Tor hyperactivation also drives rapamycin resistance in the *upf1*Δ mutant, which is suppressible by the substitution of glucose for acetate or constitutive growth at 37°C. It is tempting to speculate that NMD-mediated control of Tor may act to maintain Tor signaling in the presence of constitutive or chronic stress. An analogy for this process can be seen in *S. cerevisiae* where Tor maintains steady-state levels of pro-growth ribosome biogenesis and protein synthesis gene expression during sequential transition to less-preferred sources of carbon (86). While conservation of this mechanism in *C. neoformans* is unknown, reduction in Tor signaling in *C. neoformans* is required for thermal adaptation as overexpression of the *TOR1* kinase (*TORoe*) compromises growth at higher temperatures, which can be rescued by the addition of rapamycin (49). Growth in low-glucose conditions alters the expression of core components of Tor, but only in a mating type-dependent manner (43). A comprehensive analysis of Tor complex components and their regulation in *C. neoformans* is needed to test the model of nutrient-responsive regulation of fluconazole sensitivity and stress adaptation informed by this study.

## Methods

### Strains

*C. neoformans* var. *grubii* H99 was used as the primary WT strain in this study. The *mpk1*Δ mutant was taken from the *C. neoformans* kinase deletion set obtained through the Fungal Genetics Stock Center (87). All primers used in this study are listed in Table S1. The *upf1*Δ deletion cassette was constructed through the PCR amplification and insertion of a nourseothricin resistance cassette (NAT) flanked by ∼1000bp of homologous sequence upstream and downstream of the *UPF1* genomic locus into the pBlueScript II SK(+) (pBS). Generation of the H99 *upf1*Δ mutant was performed via biolistic transformation as previously described using the PCR-amplified *upf1*Δ::NAT cassette. The *UPF1* complementation cassette was constructed by PCR amplification of gDNA using primers located ∼1000bp upstream and downstream of the *UPF1* locus.

The amplified *UPF1* locus was then inserted into the pBS-NEO vector. An N-terminal Myc tag was incorporated into the pBS-NEO-*UPF1* vector by GenScript. The complementation cassette was amplified by PCR using the NEO-F and Upf1-comp-R primers before being transformed into the *upf1*Δ mutant strain via electroporation.

Restoration of Upf1 expression in the *upf1*Δ::*MYC-UPF1* strain was determined by western blot analysis using an anti-myc primary antibody. The *C. neoformans* KN99 WT and mutant strains used in this study were acquired from the Madhani deletion collection obtained from the Fungal Genetics Stock Center.

### Media and growth conditions

Overnight cultures were grown in 5mL of YPD (1% yeast extract, 2% peptone, and 2% dextrose) at 30°C with 250rpm shaking in snap-cap tubes. For spot plate, capsule, and melanin studies, cells were used directly from overnight cultures. For Western blotting and qPCR experiments, cells were seeded at OD_600_=0.2 in 75mL of YPD and grown in baffled flasks to mid-logarithmic (mid-log; OD_600_∼0.6) at 30°C or 37°C with 250rpm shaking prior to treatment. For Western blot experiments comparing the effects of glucose, overnight cultures were washed three times with 5mL sterile deionized water (SDW) before cells were seeded at OD_600_=0.2 in 75mL of YPD or yeast extract-peptone media (YP) supplemented with 2% sodium acetate (YPAce). For capsule induction, cells were grown in snap-cap tubes with 5mL of 0.1x Saboraud dextrose media (SD; 10% (v/v) Saboraud-dextrose media, 50mM MOPS pH=7.3). For laccase and melanin induction, cells were grown in asparagine media (ASP; 1g/L asparagine, 0.25g/L magnesium sulfate, 10mM sodium phosphate pH=6.5) or on ASP agar (ASP media, 2% agar), respectively.

### RNA-seq sample preparation and Analysis

Cultures of WT and the *upf1*Δ mutant were first grown to mid-log phase and a control sample for each strain was collected. The remaining cultures were then pelleted, resuspended in pre-warmed media, and incubated at 37°C for 1 hour before another sample was harvested. Cells were then lysed using mechanical disruption by glass beads in RLT buffer with 1% β-mercaptoethanol. RNA was then purified from lysates using the Qiagen RNeasy kit as well as the RNase-free DNase set for on-column DNase I digestion. RNA integrity was confirmed by gel electrophoresis before samples were sent for library preparation, poly(A) purification, and Illumina RNA sequencing (GENEWIZ, Azenta). Two biological replicates were sequenced for each sample.

To analyze RNA-sequencing results, FASTQ files were processed using Cutadapt (adapter trimming), STAR (alignment), and RSEM (read counting) (88–90). Cutadapt was run with a quality filter of 10 and minimum read length of 1, as well as specified to trim Illumina universal adapter sequence (AGATCGGAAGAG). STAR and RSEM were run using the FungiDB build of *C. neoformans* H99 genome (91). For differential expression analysis, read counts were subjected to a pair-wise analysis using DESeq2, outputting MA-plots and differential expression tables (92). Differential expression tables were filtered for |log_2_(foldchange)| ≥ 0.8 and a p-adjusted value ≤ 0.05. Filtered tables are provided in Data S1-4. Raw data is available under the GEO accession GSE319882. Volcano plots were generated using the EnhancedVolcano package in R. Bubble plots were generated in R using results generated using the functional annotation tool on the Database for Annotation, Visualization, and Integrated Discovery (DAVID) web service (93).

### Spot plate growth analyses

Overnight cultures were washed three times with SDW, and cell density was set to an OD_600_=1.0 for an initial 1x stock per strain. Then, five serial 10-fold dilutions were made per strain from their respective 1x stock, and 5µL of each dilution was spotted on YPD agar (YPD, 2% agar) plates supplemented with indicated drugs and incubated at the indicated temperatures. For spot plates with alternative nutrient sources, YP agar (YP, 2% agar) supplemented with 2% w/v of the indicated carbon source or Yeast nitrogen base (YNB; BD Difco) with 2% dextrose was used instead of YPD agar. Plates were photographed after three days of incubation. Plates shown are one representative of three biological replicates.

### Western blotting

Following treatment and the indicated time points, 5mL of mid-log cells were pelleted and flash frozen. Protein from thawed cells was extracted and subjected to SDS-PAGE as previously described (94). Antibodies for Phospho-eIF2α (AbCam, ab32157) and total-eIF2α (Genscript, custom) were used to probe blots as previously described (95). Detection of phospho-Mpk1 was performed using rabbit α-phospho-p44/42 MAPK antibody (Cell signaling, #4370) as previously described (53). Images shown are representatives of three biological replicates.

### Melanin and Capsule Induction

To assess melanin, overnight cultures were washed 3 times with sterile deionized water before being diluted to an OD_600_=1.0 in 10mL of sterile deionized water. These dilutions were subjected to centrifugation to pelleted cells, supernatant was decanted, and cells were resuspended in the residual supernatant to create a slurry. Then, 10µL of cell slurry was spotted onto ASP agar plates supplemented with L-dopamine (L-DOPA; 1mM) with or without the addition of various concentrations of glucose (0.1% or 1% w/v). After spots dried, the plates were then incubated at 30°C for 24 hours before being photographed.

To assess capsule induction, overnight cultures in 0.1x SD media were incubated at 37°C with 250rpm shaking. After 16-18 hours of incubation, staining with India ink was performed by mixing ink with aliquots of culture at a ratio of 1:5 by volume. Stained cells were then mounted onto glass slides before being subjected to differential interference contrast (DIC) microscopy. To quantify capsular thickness, images were first captured using a 63x DIC objective on a Leica DMi8 inverted microscope. Measurements of capsule size for 50 cells per biological replicate were performed using freehand line and measurement tools in ImageJ. Graphs represent the capsule thickness measured across all two biological replicates.

### RT-qPCR

Cells grown to mid-log phase were pelleted and resuspended in ASP media, with 5mL of cells being collected, pelleted, and flash-frozen at indicated time points. Total RNA was extracted from cells and subjected to cDNA synthesis as previously described (96) using the Applied Biosystems high-capacity cDNA reverse transcription kit (ThermoFisher). RT-qPCR for *LAC1* (CNAG_03465) and *GPD1* (CNAG_06699), using primers listed in Table S1, was performed with the qPCR SyGreen Blue mix LoROX (PCR Biosystems). Signal measurements were made on a CFX Connect real-time PCR detection system (Bio-Rad). Each experiment consisted of two technical replicates with *GPD1* serving as an internal control. Relative expression of *LAC1* was calculated using a relative standard curve method as previously mentioned (96). Data points within plots are represented by technical replicate values across 2 biological replicates.

## Acknowledgements

This work was supported by PHS grant R01 AI131977 to JCP. Tor pathway mutants are from the Madhani *Cryptococcus neoformans* (funded by R01AI100272) obtained through the Fungal Genetics Stock Center at Kansas State University.

## Supplementary Files

**Data S1**: RNA-sequencing analysis results for differentially expressed genes in WT and *upf1*Δ strains at 30°C and during a 37°C-shift.

**Table S1**: List of primers used during this study.

**Table S2**: Differentially upregulated genes in the *upf1*Δ mutant associated with utilization of various carbon and nitrogen sources.

**Table S3**: Identification of homologous Gator1 and Gator2 complex components in *C. neoformans*

## Sequencing Data Availability

Gene Expression Omnibus (GEO)

Accession GSE319882

Review token: sloxueacjxcfjkl

## Supplemental Figures and Files

**Supplementary Figure S1.**
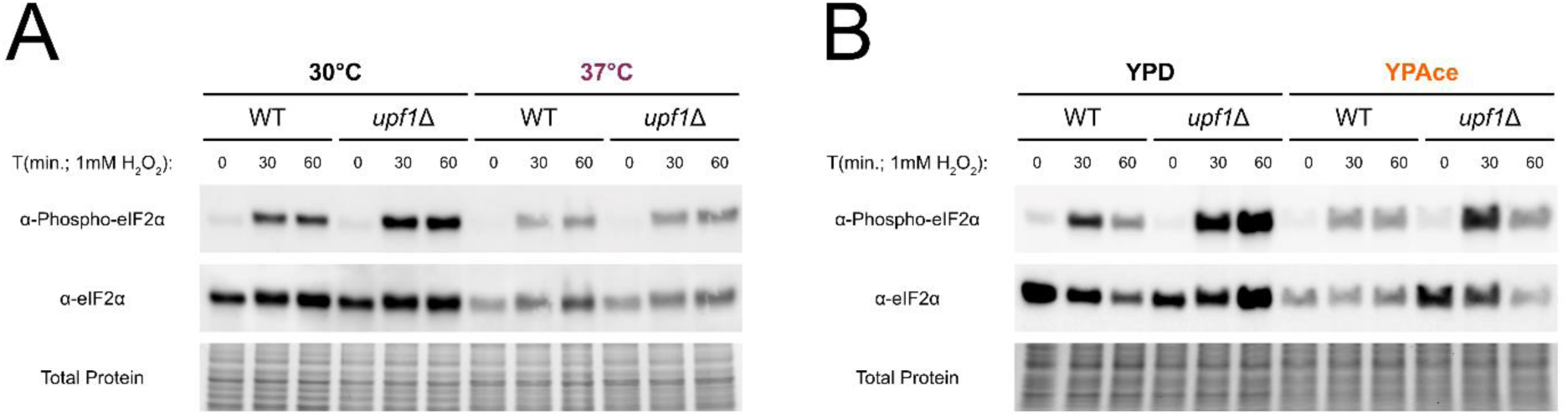
Changes in total eIF2α levels accompany changes in phospho-eIF2α in WT and *upf1*Δ strains at 37°C and in YPAce. Western blotting of phospho-eIF2α and total eIF2α in WT and *upf1*Δ strains following activation of Gcn2 by hydrogern peroxide (1mM). Protein loading was based on total protein (5µg). For time courses, only one variable was altered between experimental and control cultures per experiment, being either growth temperature (A; 30°C vs. 37°C) or growth medium (B; YPD vs. YPAce). Blots are representative of individual replicates (n=3).

